# Photocatalyst under visible light irradiation inactivates SARS-CoV-2 on an abiotic surface

**DOI:** 10.1101/2020.11.01.364364

**Authors:** Masashi Uema, Kenzo Yonemitsu, Yoshika Momose, Yoshikazu Ishii, Kazuhiro Tateda, Takao Inoue, Hiroshi Asakura

**Affiliations:** Division of Biomedical Food Research, National Institute of Health Sciences, 3-25-26 Tonomachi, Kawasaki-ku, Kawasaki, Kanagawa 210-9501, Japan; Department of Microbiology and Infectious Diseases, Toho University School of Medicine, 5-21-16 Ohmori-nishi, Ota-ku, Tokyo 143-8540, Japan; Division of Molecular Target and Gene Therapy Products, National Institute of Health Sciences, 3-25-26 Tonomachi, Kawasaki-ku, Kawasaki, Kanagawa 210-9501, Japan

**Keywords:** COVID-19, SARS-CoV-2, visible light responsive photocatalyst, RENECAT™, virucidal activity

## Abstract

There is a worldwide attempt to develop prevention strategies against SARS-CoV-2 transmission. Here we examined the effectiveness of visible light-responsive photocatalyst RENECAT™ on the inactivation of SARS-CoV-2 under different temperatures and exposure durations. The viral activation on the photocatalyst-coated glass slides decreased from 5.93±0.38 logTCID_50_/ml to 3.05±0.25 logTCID_50_/ml after exposure to visible light irradiation for 6h at 20°C. On the other hand, lighting without the photocatalyst, or the photocatalyst-coat without lighting retained viral stability. Immunoblotting and electron microscopic analyses showed the reduced amounts of spike protein on the viral surface after the photocatalyst treatment. Our data suggest a possible implication of the photocatalyst on the decontamination of the SARS-CoV-2 in indoor environments, thereby preventing indirect viral spread.

## 1. Introduction

The severe acute respiratory syndrome coronavirus 2 (SARS-CoV-2) pandemic has spread worldwide and placed countries in emerging, rapidly transforming situations. More than 44.8 million cases of COVID-19 and 1,178,475 deaths are reported to WHO as of 30 October 2020 [1]. The SARS-CoV-2 has been detected in specimens from the respiratory tract, nasopharyngeal sites, and feces in COVID-19 patients [2], The viral transmissions can occur via close human-to-human contact or via contacting a contaminated surface. To reduce the risks of environmental contamination, a myriad of disinfectants/sanitizing agents/biocidal agents are available, but their effectiveness is likely to depend on many factors such as the concentration of the agent, the reaction time, temperature, and the organic load [3]. Although the effectiveness of representative sanitizers such as ethanol and sodium hypochlorite in deactivating SARS-CoV-2 has been studied [3–5], this information is limited to a few substances and does not obtain a comprehensive picture about the effects of sanitizing agents on SARS-CoV-2.

Photocatalysts are sustainable, environmental-friendly and potent disinfectants that generate free radicals (i.e. superoxide and hydroxyl radicals) when excited by light strikes. Thus, they are efficient virucides against many pathogens including bacteria, viruses and fungi [6–7], but, to our knowledge, their effects on SARS-CoV-2 have not been investigated. In this study, we examined the virucidal activity of a photocatalyst, REN ECAT™ against SARS-CoV-2 under visible light irradiation. Our findings will contribute to the development of techniques for deactivating SARS-CoV-2 and decelerating its spread.

## 2. Materials and methods

### 2.1. Virus and cell line

The SARS-CoV-2 JPN/TY/WK-521 strain was propagated in Vero E6/TMPRSS2 cell line (JCRB 1819) [8] cultured in Dulbecco’s modified Eagle’s medium (DMEM) (Wako Pure Chemicals, Tokyo, Japan) and supplemented with 5% heat inactivated FBS (SAFC Biosciences, Lenexa, KS, USA) at 37°C in a humidified CO_2_ incubator.

### 2.2. Photocatalyst

In this study, we used the visible light responsive photocatalyst, RENECAT™ (Toshiba Materials, Kanagawa, Japan), which was mainly composed of tungsten trioxide (WO_3_). We coated 4 g/m^2^ of the photocatalyst mixed with silica binder onto 30 x 30mm soda-lime glass slides (AGC Inc., Tokyo, Japan). For the control, the glass slides were coated with silica binder alone, an auxiliary agent for the binding of the photocatalyst onto glass slides.

### 2.3. Evaluation of virucidal activity of the photocatalyst

The virucidal activity of the photocatalyst was examined following the ISO 18071 protocol [9]. Briefly, 30 μl of the virus culture medium (pH6.8) containing SARS-CoV-2 with 5% (v/v) FBS and a TCID_50_/mL of 5.93 to 6.24 log_10_ was placed on the photocatalyst-coated or a silica binder-coated (control) glass slide. Then, the slide was overlaid by a 25×25-mm VF-10 polypropylene plastic film sheet (Kokuyo Co., Ltd., Osaka, Japan) to allow a close contact between the virus and the photocatalyst. The slide was then placed in a 90-mm glass container with a wet filter paper with sterile distilled water. The container was covered with a 10×10-cm glass plate to prevent evaporation. To remove UV light with wavelength <380 nm, an acrylic resin sheet (CLAREX N-169, Nitto Jushi Kogyo Co., Ltd., Tokyo, Japan) was placed between the lamps and the samples.

The containers for the glass slides with silica binder (n=4) were placed under fluorescent lamps (FL20SSW/18, Toshiba Lighting & Technology Corporation, Kanagawa, Japan) with a luminous intensity of 3,000 lux for 6h at 20°C or 30°C (4 containers for each temperature) to select the temperature that has a lower effect on the viral titer. Moreover, to specify the incubation time, we exposed the containers without the photocatalyst (n=4 for each incubation duration) to the fluorescent light with a luminous intensity of 3,000 lux for 0h, 4h, 6h and 18h at 20°C.

To evaluate the photocatalysts virucidal acitivity, viral samples on the glass slides that were coated with (i) the photocatalyst (n=4, photocatalyst group), (ii) no photocatalyst (n=4, glass group), (iii) only silica binder (n=4, binder group) were exposed to the fluorescence lamps for 6h at 20°C. Simultaneously, four viral samples treated with the photocatalyst were kept under the dark condition (n=4, photocatalyst w/o light group). To remove UV light with wavelength <380 nm, an acrylic resin sheet (CLAREX N-169, Nitto Jushi Kogyo Co., Ltd., Tokyo, Japan) was placed between the lamps and the samples. After irradiation, the virus was recovered from glass slides by rinsing with 270 μl of serum-free DMEM, the viral suspensions and a tenfold serial dilutions were then incubated into Vero E6/TMPRSS2 cells for 3 days to calculate the TCID_50_ as described by Kärber [10]. Fig. 1A shows the experimental setup we used for this study.

**Fig 1.**
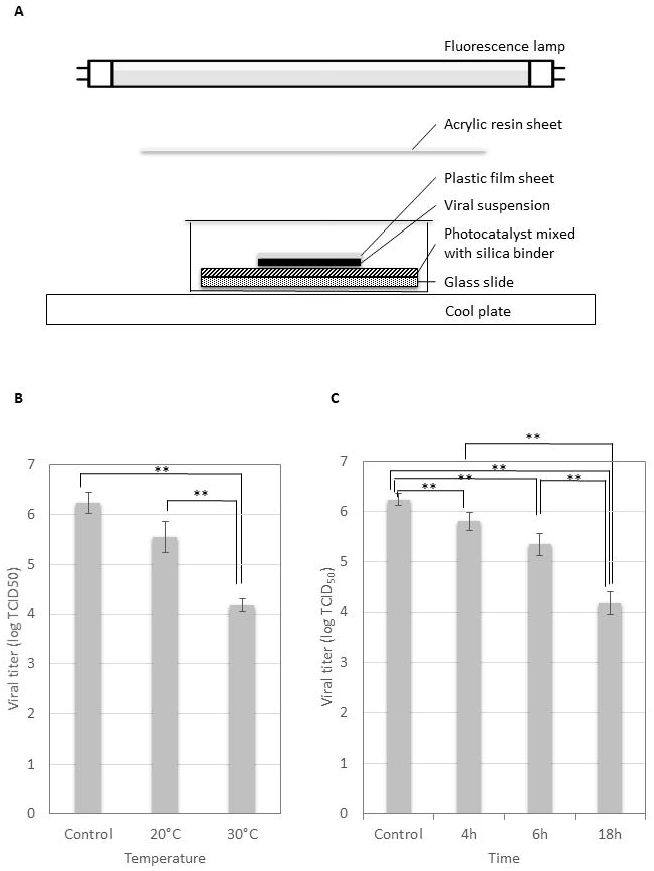
Optimization of the experimental conditions for the evaluation of virucidal activity of the photocatalyst against SARS-CoV-2. (A) Schematic representation of the experimental setup to evaluate the virucidal effects of visible light responsive WO_3_ photocatalyst. Cool plate was used to adjust the temperature on the surface of the photocatalyst. (B) Evaluation of SARS-CoV-2 stability under different temperatures (20°C, 30°C). 5% FBS-containing SARS-CoV-2 suspension was incubated on the slide glass either at 20°C or 30°C for 6h under visible light irradiation (3,000 lux). (C) Evaluation of SARS-CoV-2 stability under different incubation periods (0h, 4h, 6h, 18h). 5% FBS-containing SARS-CoV-2 suspension was incubated on the glass slide for the different periods at 20°C. Error bars indicate mean ± SD (n=4 per a group). In sections B and C, one way ANOVA was used to analyze statistical difference in the viral titer between each of the temperatures (20°C and 30°C)/ incubation periods (4h, 6h and 18h), and the control (0h). * = P < 0.05, ** = P < 0.01.

### 2.4. Immunoblot assay

The recovered viral suspensions from the photocatalyst-coated glass slides with or without the visible light for 6h at 20°C, or from the glass slide without the exposure (0h), were loaded onto 10% acrylamide gels. The proteins on the gels were transferred onto the PVDF membrane (MilliporeSigma, Burlington, MA, USA). Rabbit anti-SARS-CoV-2 spike protein antibody (#ab273074, Abcam, Cambridge, UK) and rabbit anti-SARS-CoV-2 nucleocapsid protein antibody (#GT×135357, GeneTex, Alton Pkwy Irvine, CA, USA) were used as primary antibodies. Goat HRP-conjugated anti-rabbit IgG (#4090-05, Southern biotech, Birmingham, AL, USA) was used as the secondary antibody. HRP-conjugated anti-ß-actin antibody (#HRP-60008, Proteintech, Rosemont, IL, USA) was used to detect residual actin from Vero E6/TMPRSS2 cell during preparation of viral suspension. Precision Plus protein WesternC™ pack (Bio-Rad, Hercules, CA, USA) was used for monitoring the molecular weight of the target. The protein signals were detected with the ECL detection system (Cytiva, Marlborough, MA, USA) and images were acquired with ImageQuant LAS500system (Cytiva) accordingly.

### 2.5. Electron microscopy

The recovered viral suspensions from the photocatalyst-coated glass slides with or without the visible light for 6h at 20°C, were fixed with 2.5% glutaraldehyde for 30 min at room temperature. Suspension aliquots (25 μl) were applied to 600-mesh carbon-coated copper grids, which were subjected to glow discharge. After absorption for 15 min, the grids were washed three times with water and treated for 30 s with 2% phosphotungsten acid/1% trehalose. Images were obtained using a transmission electron microscope (JOELJEM-2010).

### 2.6. Statistical analysis

One-way ANOVA test was performed with JMP 15.1 (SAS Institute, Cary, NC, USA) to analyze whether there is a significant difference in the virucidal activity between the groups. P values < 0.05 were considered statistically significant.

## 3. Results and Discussion

To our knowledge, the virucidal activity of photocatalysts against SARS-CoV-2 has not been tested. Although we followed the ISO protocol for evaluation of the photocatalyst, it remained unclear on the influences of incubation conditions (i.e. temperature and time for incubation).

Here, we examined the stability of SARS-CoV-2 on the glass slides under a variety experimental conditions including different temperatures, incubation periods, and with/without the photocatalyst. Given that temperatures >30°C reduces the stability of coronaviruses on abiotic surfaces [11], the viral stability was evaluated at 20°C and 30°C for 6 h, on the glass slide without the photocatalyst under visible light irradiation. After the 6h-incubation, the vital titer of 6.24±0.21 logTCID_50_/ml was reduced to 4.18±0.13 logTCID_50_/ml at 30°C, and to 5.55±0.31 logTCID_50_/ml at 20°C (Fig. 1B). The viral titer at 20°C was not significantly different from that of the control (0h) (P= 0.09) (Fig. 1B). Because at 30°C, the viral stability (4.18±0.14 logTCID_50_/ml) was significantly lower than the control (0h) (P< 0.01), we set up our experiments at 20°C to eliminate the effect of temperature on the viral titer while evaluating the photocatalyst’s virucidal effects.

In order to optimize the incubation periods, the viral suspensions were incubated on glass slides without the photocatalyst for 0h, 4h, 6h and 18h at 20°C under the visible light irradiation, followed by the viral titration. As shown in Fig. 1C, 18h of incubation resulted in the reduction of the viral titer to 4.18±0.22 logTCID_50_/ml, which were significantly different from that of the control (0h) (P< 0.01). The viral titer after the 4h- or 6h-incubation were relatively retained (<1.00 logTCID_50_/ml), represented as 5.81±0.18 logTCID_50_/ml or 5.35±0.22 logTCID_50_/ml, although they were significantly different from the control (0h) (P< 0.01) but no significant differences were shown between the 4h- and 6h-incubation periods (P= 0.17) (Fig. 1C). Our findings agree with a previous study showing the longer time of incubation (i.e. 24h at room temperature) reduced the viral stability [11]. Based on these data, we setup our experiments at 20°C, with the incubation time of 6h to evaluate the virucidal activity of the photocatalyst.

The experiments showed that the photocatalyst under visible light significantly reduced the titers of SARS-CoV-2, from 5.93±0.38 logTCID_50_/ml to 3.05±0.25 logTCID_50_/ml (P<0.01). The viral titer was reduced to 5.55±0.25 logTCID_50_/mL in the absence of visible light (P= 0.46) (Fig. 2A). The viral stability on the silica binder alone (with no photocatalyst; binder), and on the glass slides in the absence of photocatalyst/silica binder under florescence light (glass) were 5.18±0.13 logTCID_50_/ml and 4.88±0.13 logTCID_50_/ml, and were significantly different from that on the photocatalyst (P< 0.01 or P= 0.04), respectively (Fig. 2A). These findings indicate that the silica binder under visible light or the presence of photocatalyst substances without light did not drastically affect the viral stability while photocatalyst under visible light significantly reduced the titer of SARS-CoV-2 during the experimental periods (6h). A previous study reported that ultraviolet light (UV) at 254 nm could reduce the activity of SARS-CoV-1 [12], In this agreement, our comparative data of the photocatalyst treatment with or without visible light suggested that a part of the reduced viability of SARS-CoV-2 throughout the exposure on photocatalyst under visible light irradiation might be due to the light irradiation.

**Fig 2.**
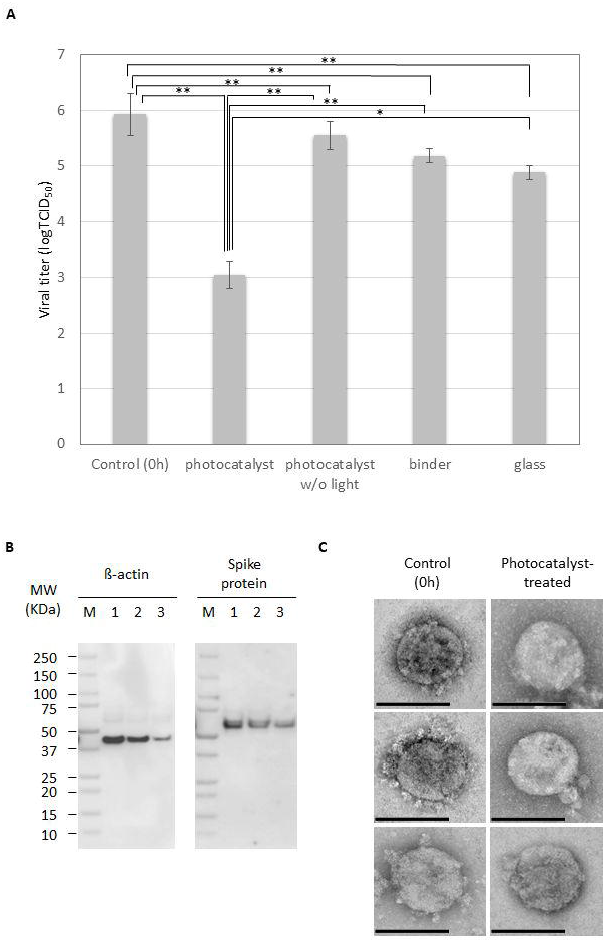
The visible light responsive photocatalyst exhibits virucidal activity against SARS-CoV-2. (A) Results of SARS-CoV-2 inactivating activity of the photocatalyst. 1% FBS-containing SARS-CoV-2 suspension on the photocatalyst was exposed to the visible light for 6h at 20°C (photocatalyst + Light) or kept under dark conditions (photocatalyst w/o Light). Also, the viral titer was measured on the silica binder (binder) or glass slide (slide alone). Error bars indicate mean ± SD (n=4 per group). ANOVA test was used to analyze statistical significance between the groups. * = P < 0.05, ** = P < 0.01. (B) Western blot analysis for the detection of the viral spike and nucleocapsid proteins as well as ß-actin protein, in the viral suspensions throughout the photocatalyst treatment. Lane M, molecular weight marker; 1, viral suspension before exposure (control); 2, viral suspension after 6h-incubation on the photocatalyst without light exposure (photocatalyst w/o light); 3, viral suspension after 6h-incubation on the photocatalyst under visible light irradiation (photocatalyst). (C) Electron micrographs of SARS-CoV-2 with or without treatment of the photocatalyst for 6h. Left panels show the images of virus treated without the photocatalyst. Right panels represent the virus treated with the photocatalyst. Bars indicate 100nm scale.

Intriguingly, immunoblot analysis showed a decreased level of the viral spike protein and residual ß-actin protein (originated from Vero E6/TMPRSS2 cells) after the 6h-treatment with the photocatalyst under visible light, compared with the sample before treatment, whereas intensity of the viral nucleocapsid protein retained even after the photocatalyst treatment (Fig. 2B). These data suggested the damage of viral surface protein and free proteins such as ß-actin by the photocatalyst treatment. In this support, electron microscopic analyses indicated the reduced amounts of spike structural molecules on the viral surface after the photocatalyst treatment (Fig. 2C). A previous report demonstrated that one of the visible light responsive photocatalysts produced free radicals such as reactive oxygen species (ROS), thereby damaging viral surface proteins of MS2 bacteriophage [13]. SARS-CoV-2 spike protein localizes on the viral surface and binds to human host cell receptor, angiotensin-converting enzyme 2 (ACE2) to establish an early course of infection [14], As recently perspective by Sun and Ostrikov [15], future study on the kinetics of affinity of the viral spike protein to human ACE2 after the photocatalyst treatment would elucidate the impact of the photocatalyst treatment on the viral infectivity. Nevertheless, our results proved evidence for the virucidal effect of the visible light responsive photocatalyst against the SARS-CoV-2, with structural damage of viral surface protein.

Disinfection technologies at indoor environments are one of the key elements in avoiding the spreads of COVID-19, particularly for medical doctors and front-line healthcare workers in hospital [16]. Most liquid biocides such as ethanol, sodium hypochlorite and electrolyzed water demonstrate prompt anti-SARS-CoV-2 activity [5,17]; however, those virucidal effects are short-term and become inactivated once contaminated with certain organic substances. By contrast, the photocatalytic techniques are expected to show relatively moderate but long-term virucidal activity because they are renewable [18]. In this study, we demonstrated the virucidal activity of the photocatalyst under a controlled experimental condition (6h-light exposure at 20°C). Several factors can affect the efficiency of photocatalyst to inactivate pathogens (i.e. pH, temperature, catalyst loading, light intensity and wavelength) [19], but our experiments were conducted under conditions to mitigate the effects of other factors, and only evaluated the virucidal activity of the WO_3_ photocatalyst against the SARS-CoV-2. In summary, our data suggest that the photocatalyst can be used to disinfect and prevent the viral spread in indoor environments in the presence of visible light. Future studies are required to clarify how long the photocatalyst could retain virucidal activity against SARS-CoV-2 throughout a continuous viral exposure. Moreover, the photocatalyst’s antiviral effects on different surfaces, such as plastic, stainless and clothes should be further investigated, because different materials affected the stability of influenza A virus differently [19] and they may have similar effects on SARS-CoV-2. Lastly, exploring viral molecules and mechanisms that are involved in the stability of SARS-CoV-2 will detect potential target molecules and pathways for deactivating this virus.

## Abbreviations

SARS-CoV-2: severe acute respiratory syndrome coronavirus 2;
TCID_50_: 50% tissue culture infective dose;
FESS: fetal bovine serum;
WHO: World Health Organization

## Acknowledgements

We wish to thank Mrs. Kayo Yoshida and Mr. Yoshihito Tsutsui from Toshiba Materials Co. Ltd. for their technical assistance during the experiments. Also, we are grateful to Dr. Masayuki Shimojima and Dr. Masayuki Saijo, from National Institute of Infectious Diseases, Japan for providing the SARS-CoV-2 JPN/TY/WK-521 strain. This study was supported by AMED under grant number 19fk0108113s0601.

The authors would like to thank Enago (www.enago.com) for the manuscript review and editing support.

